# Differences in word learning: bilingualism or linguistic experience?

**DOI:** 10.1101/551408

**Authors:** Maria Borragan, Angela de Bruin, Viktoria Havas, Ruth de Diego-Balaguer, Mila Dimitrova Vulchanova, Valentin Vulchanov, Jon Andoni Duñabeitia

## Abstract

Bilinguals may be better than monolinguals at word learning due to their increased experience with language learning. In addition, bilinguals that have languages that are orthotactically different could be more used to dissimilar orthotactic patterns. The current study examines how bilinguals with languages that are orthotactically similar and dissimilar and monolinguals learn novel words that violate or respect the orthotactic legality of the languages they know and how this learning may be affected by the similarity between the bilinguals’ two languages. In Experiment 1, three groups of children were tested: monolinguals, Spanish-Basque bilinguals (dissimilar orthotactic languages), and Spanish-Catalan bilinguals (similar orthotactic languages). After an initial word learning phase, they were tested in a recall task and a recognition task. Results showed that Spanish-Basque bilingual children performed differently than the other two groups. While Spanish monolinguals and Spanish-Catalan bilinguals recognized illegal words worse than legal words, Spanish-Basque bilinguals showed equal performance in learning illegal and legal patterns. A replication study conducted with two new groups of Spanish-Basque children (one group with high Basque proficiency and one group with a lower proficiency) indicated that the effects were not driven by the proficiency in the second language since a similar performance on legal and illegal patterns was observed in both groups. In Experiment 2, two groups of adults, monolinguals and Spanish-Basque bilinguals, were tested with the same task used in Experiment 1. The effect seen in children seems to be absent in adults. Spanish-Basque bilingual adults showed better overall learning performance than monolinguals, irrespective of the illegality of the items. Differences between groups could be due to the effect of having acquired literacy and linguistic competence.

## Introduction

Bilingualism is no longer an exceptional linguistic reality and knowing more than one language is required for business, education, and to communicate with others in many modern societies. Thus, bilingualism has become an important research area in the last decades. Despite the increasing number of studies exploring the effects of bilingualism on domain-general and domain-specific cognitive processes [1–3], the impact of bilingualism on language learning has received less attention. Previous work has suggested that bilinguals may be better at word learning than monolinguals due to their experience with language learning (see [4]). However, it is not clear whether bilinguals in general are better at word learning or whether these effects are related to and dependent on the specific characteristics of the two languages they have mastered.

Bilinguals know that objects may have different names in each of their languages and may therefore link translations in another new language more easily to a known concept than monolinguals. Along this line, studies focusing on the bilinguals’ and monolinguals’ capacity to learn a third language have suggested that bilinguals achieve a higher proficiency level in the new language than their monolingual peers [5,6]. This learning benefit has been observed both for bilinguals who learned their languages in a classroom environment [7,8], as well as for bilinguals who had acquired both languages from birth [4,9]. For instance, the latter two studies were based on a word-learning task that included novel words created to be phonologically unfamiliar to the participants. Bilingual learners had highly contrasting language combinations, such as Spanish-English or Mandarin-English. The new words had to be learned as translation equivalents of existing words from the participants’ native language. Results showed that bilinguals outperformed monolinguals in their learning performance (see [10], for review). Similar findings were also observed by Kaushanskaya and Rechtzigel [11] in a study in which English-Spanish bilinguals were contrasted with two groups of monolinguals differing in their memory span. Bilinguals outperformed both groups of monolinguals when learning novel words irrespective of the specific phonological features of the novel words and the memory span of the participants.

However, these findings do not necessarily imply that all types of bilinguals will learn novel words better than monolinguals. Werker & Byers-Heinlein [12] underscore the importance of the specific language pairs in the bilingual language system and their interaction. As seen, the abovementioned studies tested bilinguals who mastered two languages with clearly different orthotactic and phonotactic structures (e.g., Mandarin-English or Spanish-English), and it could be tentatively hypothesized that this is the underlying factor that makes the learning of new items more effective for bilinguals. Different bilingual populations speak different languages and the characteristics of the specific languages spoken may affect how known pieces of information are processed and, more importantly for the purposes of the current study, how new pieces of information are learned. Studies suggest that the structure of one’s known language(s) may determine the way new sounds are processed [13]. Furthermore, Bialystok et al. [2] demonstrated that bilinguals whose two languages share the same print-to–sound principle and/or the same writing system (i.e., Spanish-English) show better performance in a meta-phonological task (count the number of sounds in a word) than bilinguals with two languages following a different writing system (i.e., Chinese-English). Certainly, learning new phonological and orthographic patterns that also exist in one’s native language(s) is expected to be easier that learning completely different patterns (see [14]). Thus, the current study examines how bilinguals and monolinguals learn words that violate or respect the orthotactic legality of the languages they know (i.e., the language-selective pattern of grapheme combinations in written words), and how this learning may be affected by the similarity between the bilinguals’ two languages. To this end, the performance of two groups of bilinguals was compared to that of a group of monolinguals.

We hypothesized that when bilinguals have to learn new orthotactic patterns that do *not* exist in their languages, the degree of *dissimilarity* between the two languages could improve learning of these different structures or patterns. Daily management with different orthotactic patters could lead these bilinguals to be more flexible when encountering new patterns. Thus, we also conjectured that bilinguals that know languages with different orthotactics rules are more prone to accept and learn new words with different orthotactic characteristics than bilinguals with languages that are orthotactically similar.

Recent research has highlighted the critical role played by the orthotactic structure of words during bilingual visual-word recognition ([15–17], for review). Words from a given language that include certain letter combinations that are illegal in the other language known to a bilingual (namely, marked words containing language-specific orthotactic regularities) are processed differently than words whose orthotactic pattern is also plausible in the other language (namely, unmarked words; [18]). Language detection is mediated by the regularities of the sub-lexical representations of the words that are being read. Along these lines, research has demonstrated that marked words are easier to detect than unmarked words [18–20], and that they elicit lower cross-language activation levels than unmarked words [17], suggesting that language-specific orthotactic patterns represent an important cue in bilingual language processing.

With this in mind, here we investigated the extent to which monolinguals and different types of bilinguals whose language combinations critically vary in their orthotactic overlap learn new words in a different manner depending on the sub-lexical characteristics of the items. We focused on two language pairs: Spanish-Catalan and Spanish-Basque. While these three languages all share the same Roman alphabet, their sub-lexical structures vary. Spanish and Catalan share most orthotactic patterns, whereas Spanish and Basque are very dissimilar in their graphemic structure, and Basque has many bigram combinations that are illegal according to the Spanish (and Catalan) orthotactic rules.

In the first series of experiments (Experiments 1a and 1b) we will focus on children, and then the same questions will be put at test in adult samples (Experiment 2) to explore the degree of generalization across age groups and the extent to what the effects may depend on a multilingual school environment. Bilingual children attending a bilingual school need to deal with the two languages in printed materials in the same school context. They have to read in both languages and they are permanently exposed to bilingual written language. This scenario of bilingual schools where the two languages coexist are markedly different from adults’ common contexts, in which the exposure to two written languages is far less common in the same scenario. Thus, children may develop strategies different than adults to deal with this reality and its demands. Hence, we investigated if new vocabulary acquisition is easier for all types of bilinguals as compared to monolinguals (see [11]), or if this benefit depends on the specific sub-lexical characteristics of the language combination of the bilinguals, paying special attention to the orthotactic level.

Besides, we also explored whether the learning benefit of the bilinguals depends on the specific sub-lexical characteristics of the words that are being learned. To this end, we created non-existing novel orthographic representations that either respected the orthotactic structure of all the languages (e.g., the new word ‘aspilto’, which could perfectly be a word in any of the three languages according to the graphemic patterns), or that violated the orthotactic rules of these languages (e.g., the nonword ‘ubxijla’, containing the bigrams ‘bx’ and ‘jl’ that do not exist in Spanish, Catalan or Basque). We predicted that the learning benefit would be either maximal for bilinguals with more dissimilar languages at the orthotactic level on the illegal bigram combinations, since they could find it easier to deal with different orthotactic patterns due to their experience.

## Experiments 1a and 1b

### Experiment 1a

#### Methods

##### Participants

A total of 72 children (45 females; M_age_=12.9 years, SD_age_=0.8) took part in this experiment, divided into three language groups. Children were recruited from three schools located in different Autonomous Communities in Spain. First, a group of 24 Spanish monolinguals was recruited in Santander (Cantabria), which is a monolingual region located in the North of Spain. Second, a group of 24 Spanish-Catalan bilinguals was recruited in Barcelona (Catalunya), a bilingual community on the North East coast. And third, a group of 24 Spanish-Basque bilinguals was recruited in Vitoria (Basque Country), a bilingual community on the North coast.

Spanish monolinguals lived in a Spanish-only environment and attended a Spanish monolingual school. Spanish-Catalan bilingual children had acquired both languages before the age of 6. They were raised in a bilingual community and educated in a Spanish-Catalan bilingual school. Spanish-Basque bilinguals had also acquired both languages before the age of 6, and they were also attending a bilingual school. We assessed language proficiency with three different measurements (see Table 1): a subjective scale, in which participants rated their language competence on a scale from 0 to 10; a 20-item adapted version of a picture naming task [21]; the LexTale (a lexical decision task, cf., for the English version [22]; for the Spanish version [23]; and for the Basque version [21]). In addition to measuring proficiency in Spanish, Basque, and Catalan (where relevant), we also made sure that, despite English being a mandatory subject in all Spanish schools, the participants’ English level was relatively low as assessed by the English LexTale (see Table 1).

**Table 1.** Descriptive statistics of assessments.

|  | Monolinguals | Spanish-Basque<br>bilinguals | Spanish-Catalan<br>bilinguals | ANOVAs |  |
| --- | --- | --- | --- | --- | --- |
|  |  |  |  | *F(df)* | *p* |
| **Age** | 13.13 (0.90) | 12.71 (0.91) | 13.08 (0.72) | F(2,69)=1.76 | .179 |
| **Spanish competence** | 9.58 (0.97) | 9.04 (0.91) | 9.46 (0.72) | F(2,69)=2.05 | .141 |
| **Basque competence** | - | 5.38 (0.88) | - | - | - |
| **Catalan competence** | - | - | 9.25 (0.79) | - | - |
| **Spanish LexTale** | 84.44 (13.60) | 88.15 (4.87) | 82.74 (7.76) | F(2,69)=2.05 | .141 |
| **Basque LexTale** | - | 70.71 (7.03) | - | - | - |
| **Spanish picture**<br>**naming** | 99.38 (1.69) | 97.5 (2.95) | 98.13 (3.23) | F(2,69)=2.36 | .112 |
| **Basque picture naming** | - | 72.91 (2.80) | - | - | - |
| **Catalan picture naming** | - | - | 96.25 (3.69) | - | - |
| **Economic status** | 6.29 (1.12) | 6.04 (1.60) | 6.75 (0.85) | F(2,69)=2.05 | .141 |
| **IQ** | 18.17 (4.43) | 20.17 (3.45) | 20.04 (3.63) | F(2,69)=2.02 | .140 |

*Note.* Values reported are means and standard deviation in parenthesis on age (in years), subjective language competence ( 0-10 scale), LexTale (%), picture naming (% correct), economics status (1-10 scale), and IQ (correct answerers). The last column shows the results from one-way ANOVAs comparing the three language groups on the different assessments.

Participant groups were matched in age, language proficiency in Spanish, socioeconomic status, and IQ (see Table 1). Socioeconomic status was measured with a short parental questionnaire in which they were asked to indicate on a scale from 1 to 10 how they perceived their economic situation as compared to other members of their community [24]. IQ was measured with a 6-minutes abridged version of the K-BIT [25] in which participants had to complete as many matrices as they could in the time provided. As seen in Table 1, bilingual participants could not be matched on their second language competence (i.e., Basque and Catalan). Spanish-Basque bilinguals were less proficient in Basque than the Spanish-Catalan bilinguals were in Catalan, and this may be due to the origin of the Spanish-Basque bilinguals, who come from and were tested in a city in which Basque is mainly used at school, while the Spanish-Catalan participants used Catalan in daily life outside school too.

All participants were right-handed and none were diagnosed with language disorders, learning disabilities, or auditory impairments. They and their families were appropriately informed and legal guardians signed consent forms prior to the experiment. The protocol was carried out according to the guidelines approved by the BCBL Ethics Committee.

##### Materials

Thirty letter strings were created for this experiment (see Appendix 1). All strings followed the same orthographic structure of vowel, consonant bigram, vowel, consonant bigram, and final vowel (i.e., VCCVCCV). The consonantal bigram was manipulated to be legal or illegal in the three critical languages (i.e., half of them resulting in legal and half of them in illegal strings). The fifteen strings containing legal consonant bigrams (e.g., ‘ASPILTO’) included bigram combinations that were plausible in Spanish, Catalan and Basque (e.g., the consonant cluster ‘SP’ appears in ‘avispa’, the Spanish for wasp, ‘ispilu’, mirror in Basque, and ‘espai’, which corresponds to space in Catalan). The other 15 strings contained two consonant clusters that corresponded to illegal bigram combinations in all three languages (e.g., ‘UBXIJLA’, where the bigrams ‘BX’ and ‘JL’ do not exist in any of the three critical languages). The average bigram frequency of the legal words in the three languages did not statistically differ (*p*>.15; see Appendix). Each of the 30 strings was paired with a different video clip of an invented 3D object that rotated on 3 axes (see [26]). The new word stimuli were recorded in a soundproof room with a Marantz^®^ professional PMD671 recorder by a native Spanish female with neutral intonation. Legal and illegal items could be pronounced and they were fragmented in three syllables (e.g., /as/pil/to/) following the Spanish phonology, which is the common language for the three groups.

##### Procedure

Participants were individually tested during school hours. The entire experiment lasted about one hour, including the initial assessment and the two experimental phases (i.e., learning and test). All visual stimuli were presented on a 13-inch MacBook^®^ running Experiment Builder^®^ and auditory materials were presented to both ears simultaneously using Sennheiser^®^ headphones.

The experiment was divided into learning and test phases. In the learning phase, participants were first asked to learn the 30 made-up novel strings and their associated 3D invented objects. Following a fixation cross appearing for 500ms, each string-object pair was presented for 6500ms. Each 3D object was visually presented together and aligned in time with the onset of the presentation of the visual (written) and auditory representations of the corresponding novel word to show how they could sound. After the 6500ms, participants were presented with a screen requiring them to type on the keyboard the name of the object they had just learned, and they could only continue to the next trial if the string had been written correctly. Each 30 object-string association was presented three times during the learning phase, leading to 90 trials that were presented in a random order.

The testing phase included two tasks: a recall task and a recognition task. Participants first completed a recall task in which they saw each 3D object and had to write down the corresponding name that they had learned before. They were instructed to type the string that they thought corresponded to each object, even if they did not remember the whole string. After entering their responses to the objects presented in a random order, they were asked to complete a recognition task. In each of the trials of the recognition task, participants were presented with a fixation cross displayed for 500ms, immediately followed by the centered presentation of the 3D object accompanied by two response options (a correct and an incorrect string) displayed at the lower right and left sides. Incorrect responses corresponded to strings that were presented during the learning phase, but were shuffled so that they did not match the correct objects. The location of correct and incorrect options was counterbalanced across trials. Participants responded by pressing one out of two buttons on the keyboard corresponding to the location of the correct response. If no answer was given in 10000ms, the next 3D object was presented.

##### Data analysis

Firstly, a series of repeated measures ANOVAs were carried out following a 3*2 design with the factors Group (Spanish monolinguals, Spanish-Catalan bilinguals, Spanish-Basque bilinguals) and Orthotactic Structure (legal, illegal). In the recall task, we analyzed two different dependent variables. First, we analyzed the overall accuracy by considering the absolute number of correctly recalled items from the set of 30. Second, and taking into account that recall was predicted to be markedly low given the difficulty of the task and the number of items, we calculated the Levenshtein Distance (corresponding to the number of single-character substitutions, deletions, or insertions needed in each response to match the target string), and we used this as the dependent variable. A lower number of edits indicated that the response was closer to the target. In the recognition task, accuracy (percentage of errors) and reaction times (in milliseconds) were used as the dependent variables of interest. To support the absence and presence of an illegality effect in each of the language groups, we also conducted a Bayesian analysis. A Bayes factor (*BF*_10_) shows the ratio of the probability that the data were observed under the alternative hypothesis versus the null hypothesis. For instance, *BF*_10_=5 indicates that the observed data were five times more likely to have occurred under the alternative than the null hypothesis, or in the opposite way, a *BF*_10_= .2 shows that the data were more likely to be observed under the null than the alternative hypothesis. Analyses were conducted with JASP 0.8.5.

#### Results and Discussion

In the recall task, participants recalled more legal orthotactic sequences than illegal orthotactic sequences (see Table 2), *F*(1, 69)=22.93, *p*<.001, 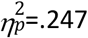. However, the main effect of Group was not significant, *F*(2, 69)=0.21, *p*=.813, 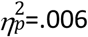., and the interaction between Group and Orthotactic Structure was not significant either, *F*(2, 69)=0.52, *p*=.594, 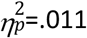. The analysis of the Levenshtein Distance showed a similar pattern, since more edits were required to transform the produced strings into the targets for illegal than for legal orthotactic sequences, *F*(1, 69)=45.41, *p*<.001, 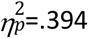. The main effect of Group, *F*(2, 69)=0.34, *p*=.713, 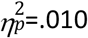, and the interaction between the two factors were not significant, *F*(2, 69)=0.35, *p*=.704, 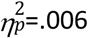.

**Table 2.** Descriptive statistics for the Recall task and the Recognition task.

|  | Monolinguals |  | Spanish-Basque bilinguals |  | Spanish-Catalan bilinguals |  |
| --- | --- | --- | --- | --- | --- | --- |
|  | Legal | Illegal | Legal | Illegal | Legal | Illegal |
| **Recall task** |  |  |  |  |  |  |
| **Recalled items** | 3.33 (5.2) | 0.28 (1.36) | 2.78 (3.89) | 0.28 (1.36) | 4.17 (7.57) | 0 (0) |
| **LD** | 5.83 (0.6) | 6.24 (0.49) | 5.76 (0.66) | 6.16 (0.38) | 5.66 (0.68) | 6.19 (0.41) |
| **Recognition task** |  |  |  |  |  |  |
| **%error** | 30.28 (16.21) | 39.72 (14.38) | 33.89 (15.59) | 33.61 (12.04) | 28.61 (18.7) | 38.33 (10.07) |
| **RT** | 1994 (628) | 2112 (716) | 1916 (661) | 2078 (843) | 1983 (566) | 1998 (651) |

Note. The table shows means and standard deviation in parenthesis on recalled items (absolute number) and Levenshtein Distance (LD) (Recall task) and %error and reaction times in ms (Recognition task) for legal and illegal orthotactic sequences for the three languages groups.

Results for the recognition task on reaction times (RT) showed that illegal orthotactic sequences tended to need more time to be responded than legal ones (see table 2), however this effect was not significant, *F*(1, 69)=2.90, *p*=.093, 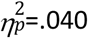. Also, the main effect of Group was not significant, *F*(2, 69)=0.07, *p*=.932, 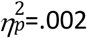, and the interaction between Orthotactic structure and Group was not significant either, *F*(2, 69)=0.57, *p*=.567, 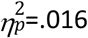. This means that all groups invested the same amount of time in all responses, with a slight tendency towards greater effort for illegal sequences. In terms of accuracy, there was a significant main effect of orthotactic sequences (see Table 2), *F*(1, 69)=12.85, *p*<.001, 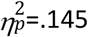. Overall, participants were more accurate at recognizing the correct word for the object when it was a legal orthotactic sequence than an illegal one. On the other hand, the main effect of Group was not significant, *F*(2, 69)=0.098, *p*=.907, 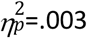, but the interaction between the two factors was significant, *F*(2, 69)=3.51, *p* = .035, 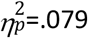. This interaction suggests that the illegality effect differs between the three groups. We therefore assessed this effect in each group separately. Spanish-Catalan bilinguals (*t*(23)=3.10, *p* =.005, Cohen’s *d*=.633, *BF*_10_=8.68) and monolinguals (*t*(23)=2.97, *p* =.007, Cohen’s *d*=.606, *BF*_10_=6.62) showed a significant effect of illegality. In contrast, this effect was not observed for Spanish-Basque bilinguals (*t*(23)=0.099, *p*=.922, Cohen’s *d*=.020, *BF*_10_ =0.21), showing that they had learned illegal orthotactic sequences to the same extent as legal ones (see Fig 1). To follow up on this interaction, we also looked at the simple main effects of Group on each level of Orthotactic Structure (i.e., on legal and illegal patterns separately). In a one-way ANOVA we found no significant effect of group for the legal (*F*(2, 69=.61, *p*=.545, 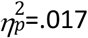) or the illegal orthotactic sequences, (*F*(2, 69)=1.63, *p*=.203, 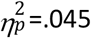). This means that the interaction between group and orthotactic sequence is not driven by the Spanish-Basque bilinguals performing better on the illegal sequences nor doing worse on the legal ones, simply that they perform the same on legal and illegal patterns whereas the other language groups perform worse on the legal than the illegal sequences.

**Fig 1.**
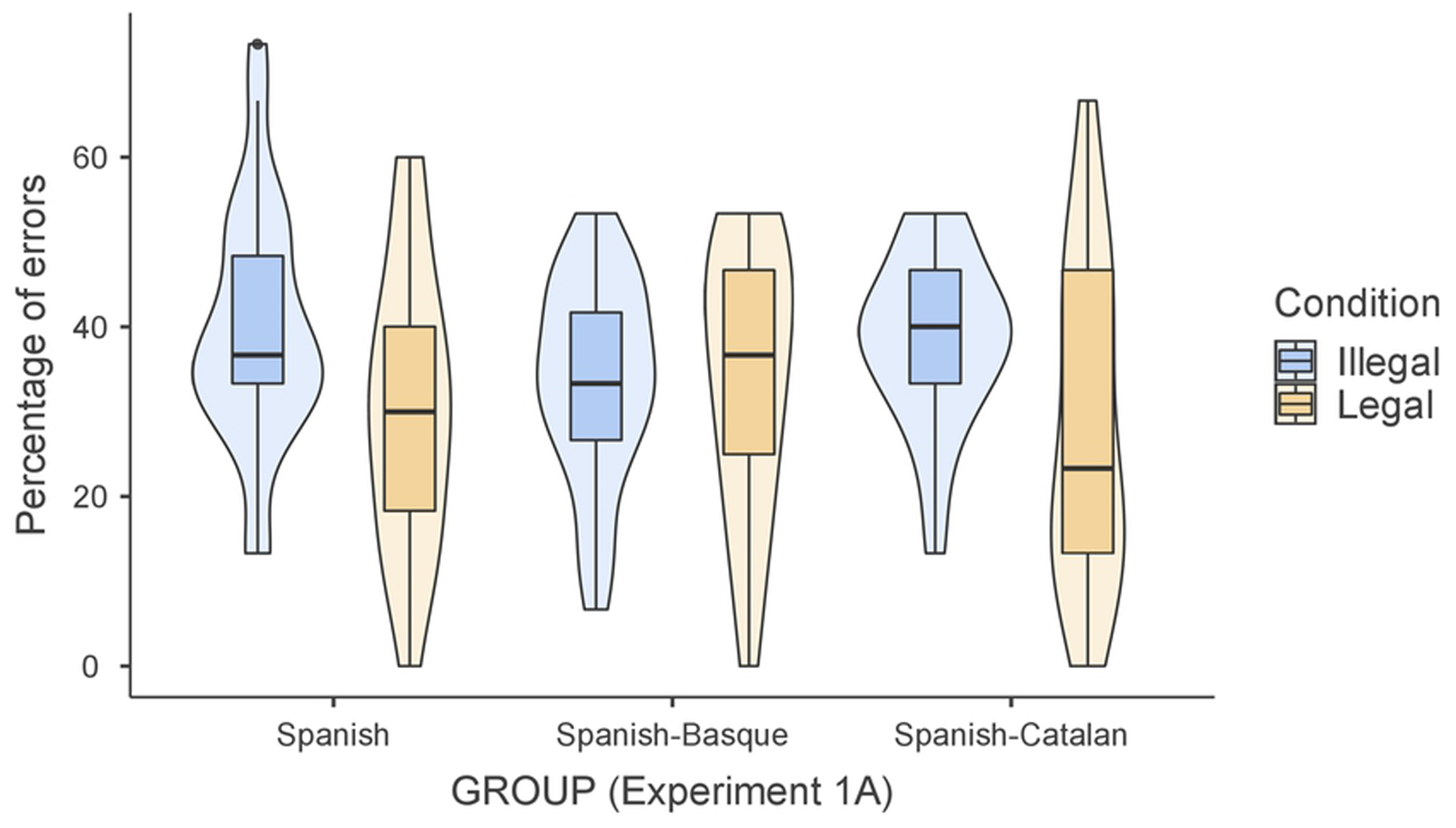
Violin plots of the percentage of errors during the recognition task for legal and illegal orthotactic sequences for each language group.

Experiment 1a aimed to examine if and how bilingual children’s linguistic experience affects the way they learn new words that violate or respect the orthotactic patterns of the languages they know. We therefore compared monolingual children’s performance to that of two groups of bilinguals: one group of (Spanish-Catalan) bilinguals who speak two languages with similar orthotactic patterns and one group (Spanish-Basque) speaking two languages that have different orthotactic patterns. Results in the recall task showed that legal words were remembered better than illegal words, but no differences between the three language groups were found in this regard. However, the recognition task showed an interaction between language group and illegality on accuracy, suggesting that monolinguals, Spanish-Catalan bilinguals, and Spanish-Basque bilinguals differ in the way they learnt new legal and illegal sequences. While monolinguals and Spanish-Catalan bilinguals recognized illegal sequences worse than the legal ones, Basque-Spanish bilinguals did not show this effect. This result suggests that group differences in word learning are not due to bilingualism as such but rather related to the two specific languages that they know. Spanish and Basque are more dissimilar (e.g., in grammar, letter sequences, phonology) than Spanish and Catalan. Therefore, the absence of a legality effect in the Spanish-Basque bilinguals could be due to their linguistic experience with the two distinct languages and the process of literacy acquisition (having already acquired the two languages).

In the next experiment (Experiment 1b), we firstly wanted to replicate the null result of illegality in Spanish-Basque bilinguals. Furthermore, as can be seen in Table 1, Basque proficiency in the group of Spanish-Basque bilinguals was lower than the Catalan proficiency in the Spanish-Catalan bilinguals. For this reason, we included two groups of Spanish-Basque bilinguals in Experiment 1b: One similar to the previous study and one group with a higher Basque proficiency. If the absence of an illegality effect is only found in the group of Spanish-Basque bilinguals with a lower Basque proficiency level, the effect in Experiment 1a may be driven by proficiency differences between the two bilingual groups. In contrast, if we do not observe an illegality effect in either group of Basque speakers in Experiment 1b, this would support our interpretation that the findings in Experiment 1a are related to linguistic experience.

### Experiment 1b

#### Methods

##### Participants

Forty-six Spanish-Basque bilingual children took part in this experiment (34 females; M_age_=12.9 years, SD_age_=0.6). Participants were recruited from two different Basque communities in the Basque Country. The first group of participants consisted of 22 Spanish-Basque bilinguals from Donostia-San Sebastian, a dense bilingual environment. The other group was composed of 24 Spanish-Basque bilinguals from Vitoria-Gasteiz, as in Experiment 1a.

Participants were matched on their language proficiency in Spanish and English, their socioeconomic status, and their IQ, as in Experiment 1a (see Table 3). However, the two Basque groups differed in their subjective measure of competence in Basque and their picture-naming performance in Basque (see Table 3).

**Table 3.** Descriptive statistics of assessments

|  | Highly proficient Basque | Less proficient Basque | T-test |  |
| --- | --- | --- | --- | --- |
|  | bilinguals | bilinguals | t(df) | p |
| Age | 13.05 (0.72) | 12.79 (0.59) | t(44)=1.31 | .197 |
| Spanish competence | 9.5 (0.86) | 9.21 (0.59) | t(44)=1.35 | .183 |
| Basque competence | 7.68 (1.09) | 5.71 (1.37) | t(44)=5.38 | <.001 |
| Spanish Lextale | 85.87 (5.59) | 87.05 (5.17) | $t(44)=0.74$ | .462 |
| Basque Lextale | 69.82 (7.49) | 71.21 (8.60) | $t(44)=0.58$ | .563 |
| Spanish picture naming | 87.73 (27.11) | 97.71 (4.66) | $t(44)=0.34$ | .729 |
| Basque picture naming | 77.45 (2.69) | 67.83 (2.45) | $t(44)=3.11$ | .003 |
| Socioeconomic status | 6.55 (1.14) | 6.25 (1.03) | $t(44)=0.92$ | .362 |
| IQ | 18.73 (2.12) | 18.38 (3.03) | $t(44)=0.45$ | .653 |

*Note.* Values reported are means and standard deviation in parenthesis on age (in years), subjective language competence (0-10 scale), LexTale (%), picture naming (% correct), socioeconomic status (1-10 scale), and IQ (number of correct answers in the timed test). The last column shows the results from the t-test comparing the two Spanish-Basque groups on the different assessments.

As in Experiment 1a, all participants’ parents received an information letter and a parental written informed consent, which was signed and returned before testing. The study was approved by the BCBL Ethics Committee. None of the children were left-handed and none were diagnosed with language disorders, learning disabilities, or auditory impairments.

##### Materials and Procedure

Materials and procedure were identical to those used in Experiment 1a.

#### Results and Discussion

We performed repeated measures ANOVAs with Group (highly proficient Basque bilinguals and less proficient Basque bilinguals) and Orthotactic Structure (legal, illegal) on accuracy and Levenshtein Distance in the recall task and percentage of error and reaction times in the recognition task. In the recall task, participants recalled more legal than illegal words (see Table 4), *F*(1, 44)=13.57, *p*=<.001, 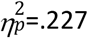, and the items recalled were closer to the target item in the case of the legal as compared to the illegal words when the Levenshtein Distance was taken into account (see Table 4), *F*(1, 44)=26.97, *p*=<.001, 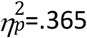. However, there were no effects of Group on accuracy, *F*(1, 44)=1.18, *p*=.282, 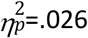, or Levenshtein distance of the recalled item, *F*(1, 44)=2.49, *p*=.122, 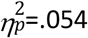. There was also no interaction between the illegality effect and Group on accuracy, *F*(1, 44)=2.09, *p*=.155, 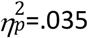, or Levenshtein distance, *F*(1, 44)=2.95, *p*=.093, 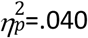. In the recognition task, participants required more time to recognize illegal words than legal ones, *F*(1, 44)=11.78, *p*=<.001, 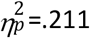, but no differences between groups were observed, *F*(1, 44)=1.12, *p*=.296, 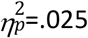, nor an interaction, *F*(1, 44)=0.11, *p*=.742, 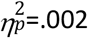. Nevertheless, we observed that participants recognized legal and illegal words equally in terms of percentages of errors, *F*(1, 44)=2.19, *p*=.146, 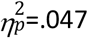, and no differences between groups were found, *F*(1, 44)=0.19, *p*=.665, 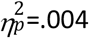 nor an interaction, *F*(1, 44)=0.15, *p*=.699, 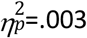, showing that the lack of illegality effect was similar for both groups of Spanish-Basque bilinguals (see Fig 2).

**Table 4.** Descriptive statistics for the Recall task and the Recognition task.

|  | High proficient Basque bilinguals |  | Less proficient Basque bilinguals |  |
| --- | --- | --- | --- | --- |
|  | Legal | Illegal | Legal | Illegal |
| **Recall task** |  |  |  |  |
| Recalled items | 3.03 (4.47) | 1.21 (2.63) | 5.56 (6.42) | 1.39 (5.55) |
| LD | 5.8 (0.50) | 6.09 (0.43) | 5.38 (0.92) | 5.95 (0.69) |
| **Recognition task** |  |  |  |  |
| %error | 30.61 (12.46) | 33.94 (12.83) | 29.72 (16.68) | 31.67 (12.00) |
| RT | 1959 (637) | 2153 (785) | 1755 (505) | 1991 (546) |

Note. Mean and standard deviation in parenthesis on recalled items (absolute number) and Levenshtein Distance (LD) (Recall task) and % of errors and reaction times in ms (Recognition task) for legal and illegal orthotactic sequences for the three language groups.

**Fig 2.**
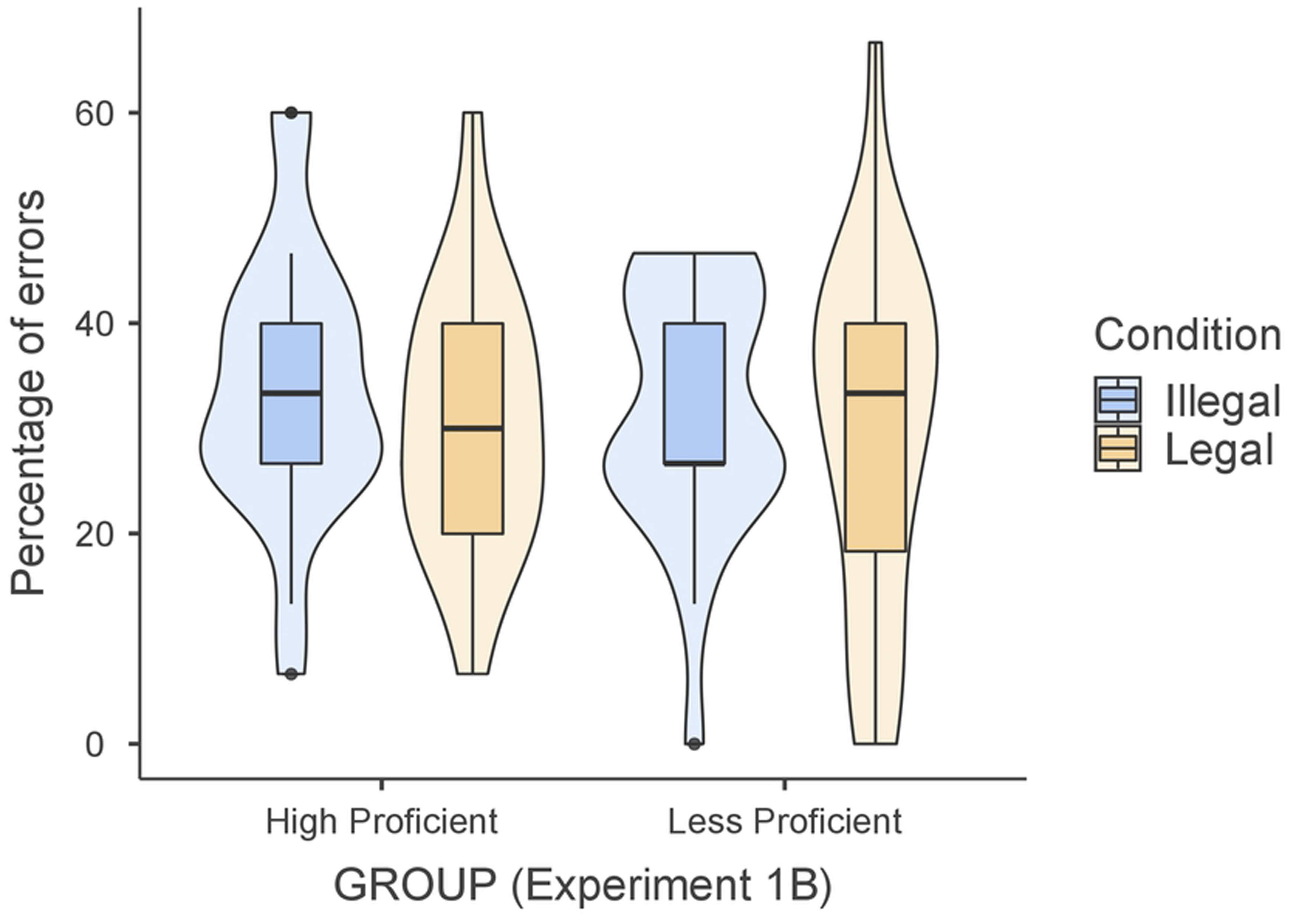
Violin plots of the percentage of errors in legal and illegal orthotactic sequences.

We investigated whether the effects were due to the characteristics of the languages or the proficiency of the children. Thus, Experiment 1b aimed to replicate the findings from the Spanish-Basque bilingual children tested in Experiment 1a in two new samples of Spanish-Basque bilinguals (a group of more balanced bilinguals and a group with the same proficiency as in Experiment 1). Similar to Experiment 1a, these bilingual children recalled more legal than illegal words, and importantly, they recognized legal and illegal words to the same extent. Furthermore, no differences were observed between these two groups regardless of their proficiency differences, suggesting that the (absence of an) illegality effect was not modulated by proficiency in Basque. Thus, these findings provide support to the results from Experiment 1a, suggesting that linguistic experience with languages that differ from each other at the orthotactic level may modulate word learning in bilingual children.

## Experiment 2

Next, we wanted to test whether the same differential pattern observed in Spanish-Basque bilingual children is also present in Spanish-Basque bilingual adults, or whether there is a different developmental trajectory of these effects that make the absence of sensitivity to orthographic markedness during new word learning (e.g. flexibility in word learning) diminish as a function of age. We therefore conducted a study using the same methodology as in Experiment 1 comparing Spanish-Basque bilingual adults to Spanish monolingual adults. If experience speaking two different languages affects learning new legal versus illegal words across the lifespan, we would expect patterns similar to those reported in Experiment 1 (i.e., Spanish-Basque bilinguals recognizing illegal and legal words to the same extent and Spanish monolinguals remembering legal words better than illegal ones). However, considering that children are still in the process of acquiring and conforming the repertoires of their two languages, if the findings observed in Experiment 1 are related to this ongoing language development and acquisition of vocabulary and literacy, we may observe different patterns in adults.

### Methods

#### Participants

Forty-eight adults took part in this experiment (30 females; M_age_=21.68 years, SD_age_=2.8), twenty-four were Spanish monolinguals from the University of Cantabria and the other twenty-four were Spanish-Basque bilinguals from the University of the Basque Country. As in Experiment 1, they were matched on their language proficiency in Spanish and English, socioeconomic status, and IQ (see Table 5).

**Table 5.**
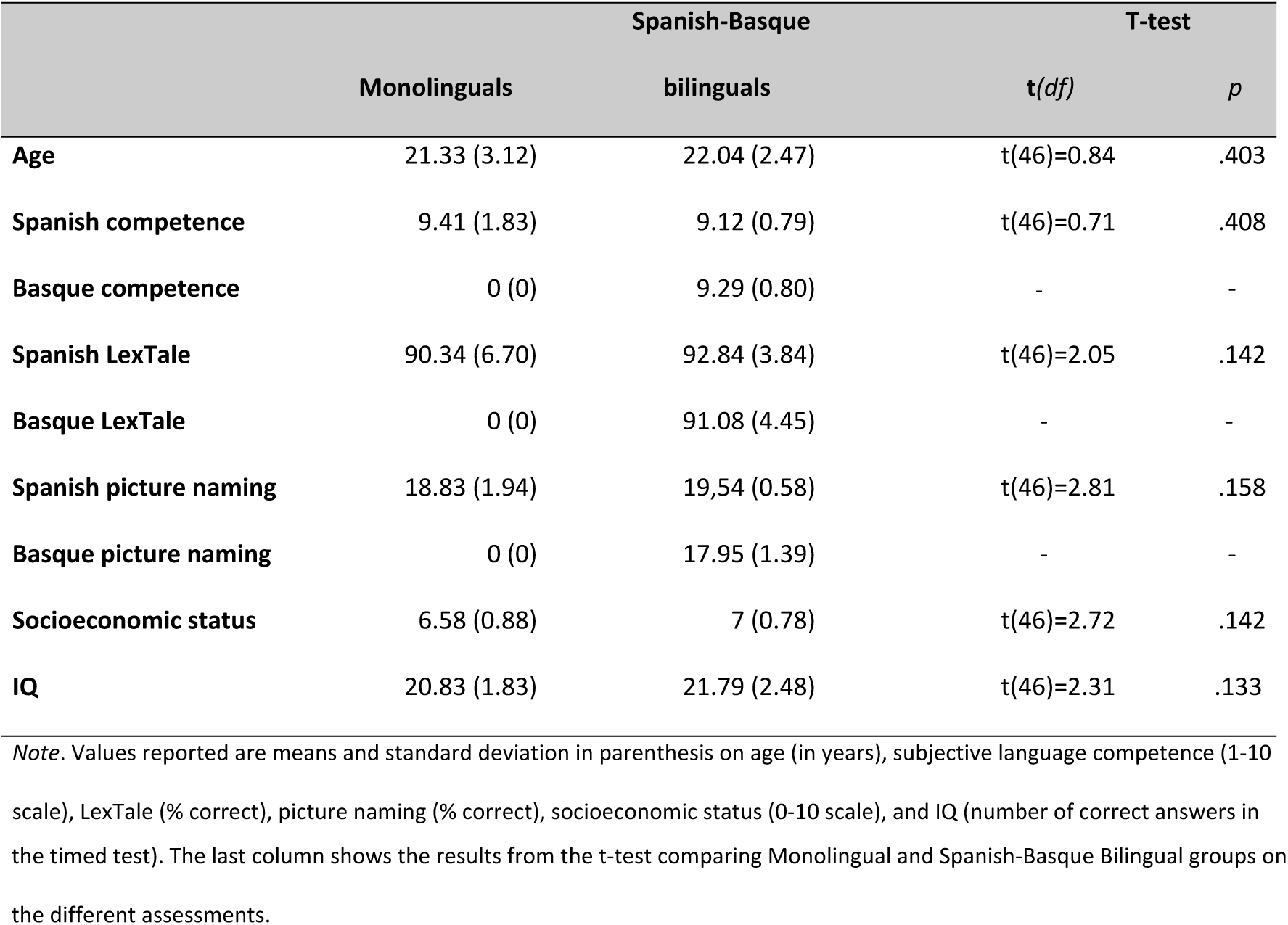
Descriptive statistics of assessments.

The study was approved by the BCBL Ethics Committee and all participants signed an informed consent before the experiment and were compensated for their time. None of them were left-handed and none were diagnosed with language disorders, learning disabilities, or auditory impairments.

#### Materials and Procedure

Materials and procedure were identical to those used in Experiment 1.

## Results and Discussion

Repeated measures ANOVAs were conducted with the factors Group (Spanish monolinguals, Spanish-Basque bilinguals) and Orthotactic Structure (legal, illegal) on accuracy and Levenshtein distance in the recall task and percentage of errors and reaction times in the recognition task. In the recall task, participants recalled more legal than illegal orthotactic sequences, *F*(1, 46)=16.92, *p*=<.001, 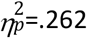, but groups did not differ on performance, *F*(1, 46)=0.47, *p*=.492, 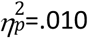, and there was no interaction between Group and Orthotactic Structure, *F*(1, 46)=1.57, *p*=.217, 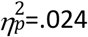. The analysis of Levenshtein Distance showed that the recall of legal orthotactic sequences was closer to the template than that of illegal sequences, *F*(1, 46)=51.97, *p*=<.001, 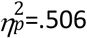. Also, a Group effect was found, *F*(1, 46)=11.54, *p*=.001, 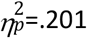, such that the Spanish-Basque bilinguals outperformed the monolinguals. The interaction between the two factors was significant, *F*(1, 46)=4.83, *p*=.033, 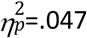, suggesting that Spanish-Basque bilingual adults’ recall was more similar to the target items, especially in the case of the legal sequences (see Table 6).

**Table 6.** Descriptive statistics for the Recall task and the Recognition task

|  | Monolinguals |  | Spanish-Basque bilinguals |  |
| --- | --- | --- | --- | --- |
|  | Legal | Illegal | Legal | Illegal |
| **Recall task** |  |  |  |  |
| **Recalled items** | 1.04 (1.90) | 0.38 (1.44) | 1.63 (2.26) | 0.38 (0.58) |
| **LD** | 5.69 (1.09) | 6.18 (0.86) | 4.58 (1.18) | 5.51 (0.66) |
| **Recognition task** |  |  |  |  |
| **%error** | 25.83 (12.64) | 35.83 (19.14) | 11.11 (13.43) | 24.72 (11.37) |
| **RT** | 1974 (444) | 2256 (637) | 1899 (433) | 2318 (642) |

*Note.* Mean and standard deviation in parenthesis on recalled items (absolute number) and Levenshtein Distance (LD) (Recall task) and %error and reaction times in ms (Recognition task) for legal and illegal orthotactic sequences for the three language groups.

In the recognition task, adults showed a main effect of Orthotactic Structure on the percentage of errors, *F*(1, 46)=31.05, *p*=<.001, 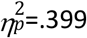, as well as of Group, *F*(1, 46)=12.91, *p*=<.001, 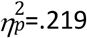, but not an interaction, *F*(1, 46)=0.73, *p*=.399, 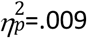. This means that adults recognized the legal orthotactic structures more accurately than the illegal ones, and that Spanish-Basque bilinguals outperformed monolinguals in the overall learning (see Fig 3). However, the absence of interaction suggested that both language groups remembered the legal words better than the illegal ones to a similar extent. The analysis of the reaction times in this recognition task showed that adults recognized legal orthotactic sequences faster than illegal ones, *F*(1, 46)=28.14, *p*=<.001, 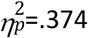. However, there was no Group effect, *F*(1,46)=0.002, *p*=.965, 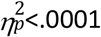, nor an interaction, *F*(1, 46)=1.09, *p*=.303, 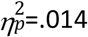.

**Fig 3.**
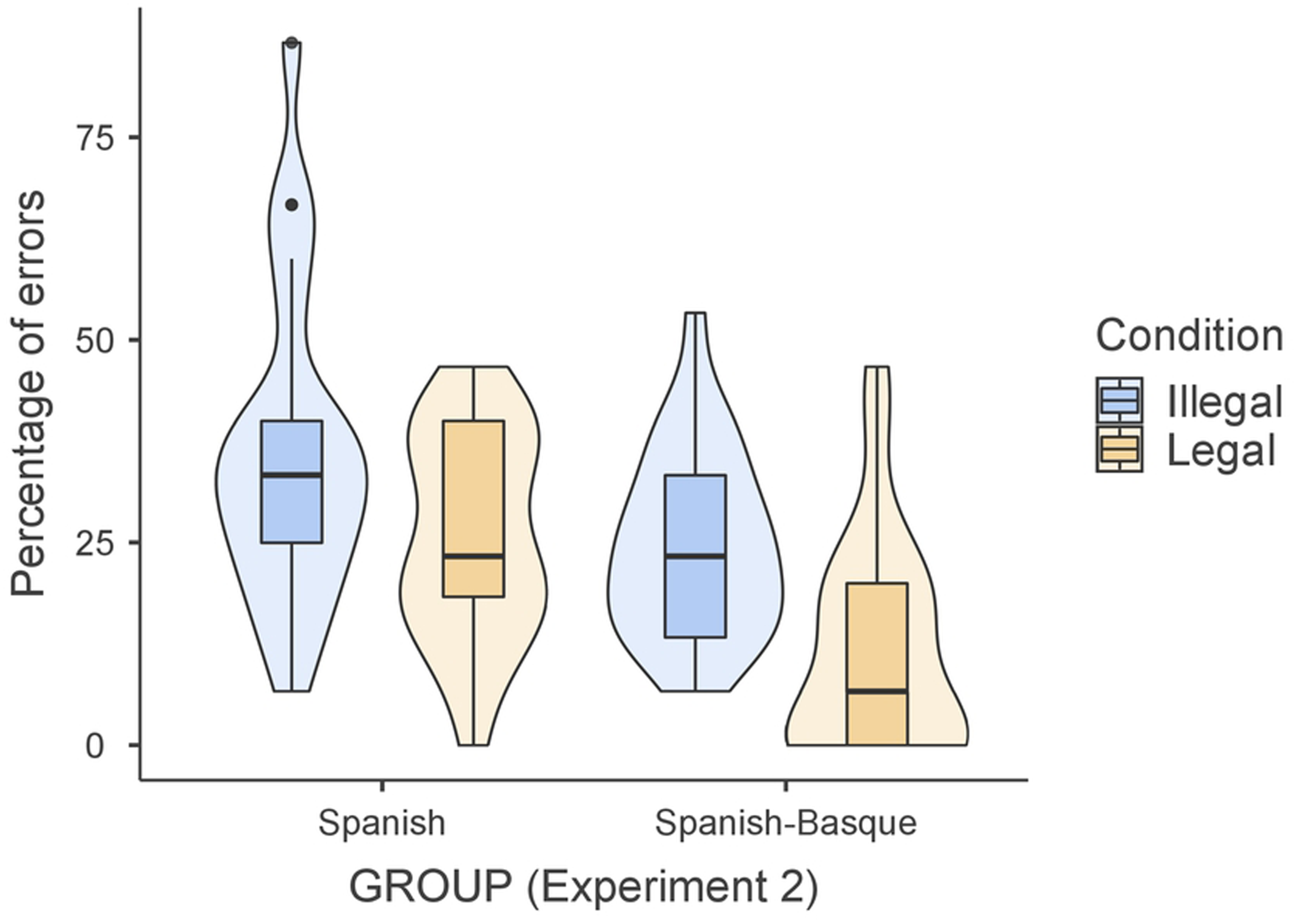
Violin plots of the percentage of errors in legal and illegal orthotactic sequences.

Experiment 2 assessed whether Spanish-Basque bilingual adults differed from Spanish monolingual adults, as children did, in the way they learn new legal and illegal words. Similar to the results reported in Experiments 1a and 1b, the recall task showed that both language groups recalled more legal than illegal words. However, unlike in Experiment 1, the recognition task showed a legality effect for both language groups. Both the Spanish monolinguals and the Spanish-Basque bilinguals remembered legal words better than illegal words. In terms of overall performance, Spanish-Basque bilinguals outperformed Spanish monolinguals: bilinguals remembered more words than monolinguals in the recognition task and their responses in the recall task were closer to the target words.

These results are in line with the literature showing an advantage in word learning in bilinguals compared with monolinguals (for a review, see [27]), aligning with previous studies focused on adult word learning showing that bilinguals outperform monolinguals [4,9,11,28]. The results from Experiment 2 provide support to this view, suggesting that bilinguals can benefit from their previous experience with language (e.g., learning vocabulary in various languages) to achieve a higher level of performance in new word learning tasks. However, their experience with two languages differing in terms of orthotactic structures did not affect performance.

## General Discussion

Previous research suggests that bilinguals may be more efficient than monolinguals at word learning due to their experience with language learning [4,9]. The aim of the present study was to examine whether new word learning is driven by the bilingual experience itself, or rather the specific linguistic experience with the languages in the bilingual pair. We were specifically interested in whether greater language difference can give an edge in novel word learning. Therefore, we conducted two experiments to test this hypothesis. In Experiment 1, we asked children to learn new words containing legal or illegal patterns. Specifically, in Experiment 1a we tested children that were Spanish-Basque bilinguals, Catalan-Spanish bilinguals, or Spanish monolinguals and in Experiment 1b we tested two additional groups of Spanish-Basque bilinguals in an attempt to replicate the findings and control for the effects of proficiency. In a second experiment, we carried out the same task as in Experiment 1, but with two groups of adults (Spanish monolinguals and Spanish-Basque bilinguals) in order to test whether the effects were only present in the process of language development.

In Experiment 1a, we observed that the children in all three language groups recalled legal items better than illegal ones, in line with prior literature showing that it is easier to learn items that correspond with our prior knowledge [14]. No effects of language group were observed. However, this recall task is not very informative due to the low percentage of words the children were able to properly recall. This low performance could lead to a floor effect, in which we cannot observe a difference between groups simply because performance does not allow for enough variability [29]. A greater amount of rehearsal might be needed to enhance recall performance and show further effects of language group. Due to the difficulty of the task, recognition memory may be more sensitive to showing the nuanced effects of language group. Indeed, in this task Spanish-Catalan bilinguals and Spanish monolingual children showed a benefit for legal items, whereas Spanish-Basque bilingual children did not. In other words, Spanish-Basque bilingual children did not show a legality effect, and they recognized legal and illegal (namely, orthographically unmarked and marked) strings similarly. Importantly, the results of Experiment 1b with two additional groups of Spanish-Basque bilingual children demonstrated that the absence of a legality effect in this population is a stable phenomenon that does not depend on the level of proficiency.

Focusing on the language group differences in the recognition task, both monolinguals and Spanish-Catalan bilinguals follow the patterns of learning that are described in prior literature [14]—namely, they learn legal sequences better than illegal ones—whereas the Spanish-Basque bilinguals deviate from this. Our hypothesis is that the driving factor leading to this differential effect could be linguistic experience, meaning that by learning (or knowing) two languages that differ very strongly in their orthotactic rules, these bilinguals are less affected by the legality of new words. That is, Spanish-Basque children could be less sensitive, and thus more flexible, to the legality of words due to their experience with languages that already have patterns that violate the rules in the other language. In particular, Spanish and Basque differ strongly in their orthotactics rules, which is not so much the case with Spanish and Catalan. This experience of managing two different sets of rules is what sets this group of Spanish-Basque bilinguals apart and may have allowed them to learn words equally regardless of whether their orthotactic patterns violated the rules in the already known languages.

One important follow-up question was whether this pattern of results is only found in childhood, when language development is still ongoing, or is maintained through adulthood. In an attempt to respond to this question, in Experiment 2 we tested Spanish monolingual and Spanish-Basque bilingual adults with the same materials. Results showed a different pattern of word learning as compared to that of children, with Spanish-Basque bilingual adults clearly outperforming the monolinguals overall, but with both language groups showing a comparable legality effect. That is, contrary to the children, both the Spanish monolingual and the Spanish-Basque bilingual adults recognised legal words better than illegal words, and the former group had an overall poorer performance than the latter one.

While it is unclear what the underlying reason is for these differences between children and adults, we tentatively suggest that they are due to already acquired linguistic competence in the adults and the still ongoing language and cognitive development in children. The ability to learn a new language changes across the lifespan and children may acquire new languages and new vocabulary in a different manner than adults [30]. Children that are still learning a language, learn more vocabulary and have a greater amount of exposure to new language elements (e.g., rules, vocabulary, complex sentences) than adults [31]. In contrast, adults undergo fewer changes in the language development of the languages they already speak. Thus, considering that children can be more flexible at learning new words than adults, one could tentatively account for the fact that only Spanish-Basque bilingual children, but not adults, learn legal and illegal new words equally well. The group of children were still in the process of conforming to the Basque and Spanish vocabulary, a process that could have made them less sensitive to illegality. On the other hand, adults who have already fully acquired both languages may have also developed sensitivity to differences between languages, something that they can use as a strategy in some circumstances (see [19] for a review). Thus, contrary to children, Spanish-Basque adults may be sensitive to legal versus illegal orthotactic patterns and may consequently show better performance for legal words.

In sum, having experience with languages that differ at the orthographic (or orthotactic), but also phonotactic, level can affect word learning. Bilingual children who are exposed to two languages that have clearly different orthotactic regularities and immersed in a school context with a strong presence of written text in both languages, perform differently on word learning tasks as compared to other bilingual or monolingual children, providing them with a specific form of learning flexibility with respect to orthographic markedness. Further studies should try to disentangle the immediate causes and limitations of this phenomenon, particularly throughout the lifespan.

## Acknowledgments

This research has been partially funded by grant PSI2015-65689-P from the Spanish Government to JAD, by the AThEME project funded by the European Union (grant number 613465), by a personal grant from La Fundación La Caixa ID 100010434 to MB (code LCF/BQ/ES16/11570003), and by grant Centro de Excelencia Severo Ochoa SEV-2015-0490 by the Spanish Government. The funders had no role in study design, data collection and analysis, decision to publish, or preparation of the manuscript.

## Author contributions

Conceived the idea: MB AdB JAD RdD MDV VV VH. Designed the experiments: MB JAD AdB. Collected the data: MB. Analyzed the data: MB AdB JAD. Drafted the paper: MB under the supervision of AdB and JAD. Discussed the findings and revised the manuscript: MB AdB JAD RdD MDV VV VH.

## Supporting Information

S1 Appendix. **Thirty new words with their average bigram frequency (appearance per million). Bigram frequency is calculated averaging the frequencies of the critical consonantal bigrams**. The items are numbered in the same order as in S2 Appendix.

(PDF)

